# Machine learning applied to whole-blood RNA-sequencing data uncovers distinct subsets of patients with systemic lupus erythematosus

**DOI:** 10.1101/647719

**Authors:** William A Figgett, Katherine Monaghan, Milica Ng, Monther Alhamdoosh, Eugene Maraskovsky, Nicholas J Wilson, Alberta Y Hoi, Eric F Morand, Fabienne Mackay

## Abstract

**Objective:** Systemic lupus erythematosus (SLE) is a heterogeneous autoimmune disease that is difficult to treat. There is currently no optimal stratification of patients with SLE, and thus responses to available treatments are unpredictable. Here, we developed a new stratification scheme for patients with SLE, based on the whole-blood transcriptomes of patients with SLE.

**Methods:** We applied machine learning approaches to RNA-sequencing (RNA-seq) datasets to stratify patients with SLE into four distinct clusters based on their gene expression profiles. A meta-analysis on two recently published whole-blood RNA-seq datasets was carried out and an additional similar dataset of 30 patients with SLE and 29 healthy donors was contributed in this research; 141 patients with SLE and 51 healthy donors were analysed in total.

**Results:** Examination of SLE clusters, as opposed to unstratified SLE patients, revealed underappreciated differences in the pattern of expression of disease-related genes relative to clinical presentation. Moreover, gene signatures correlated to flare activity were successfully identified.

**Conclusion:** Given that disease heterogeneity has confounded research studies and clinical trials, our approach addresses current unmet medical needs and provides a greater understanding of SLE heterogeneity in humans. Stratification of patients based on gene expression signatures may be a valuable strategy to harness disease heterogeneity and identify patient populations that may be at an increased risk of disease symptoms. Further, this approach can be used to understand the variability in responsiveness to therapeutics, thereby improving the design of clinical trials and advancing personalised therapy.

## INTRODUCTION

Systemic lupus erythematosus (SLE) is a debilitating chronic autoimmune condition characterised by the activation of inflammatory immune cells and the production of pro-inflammatory autoantibodies responsible for pathology in multiple organs.^1^ SLE is highly heterogeneous, and can be seen as a syndrome rather than a single disease.^2^ The responsiveness of patients to available treatments is variable and difficult to predict. Rather than a small number of highly associated loci, over 60 SLE low-association loci have been identified by genome-wide association studies.^3–6^ SLE has been studied using numerous useful mouse models, each of which manifest SLE-like symptoms underpinned by different molecular mechanisms. Two examples are mice overexpressing B cell activating factor of the TNF family (BAFF, also known as TNFSF13B) i.e. BAFF-transgenic mice, in which low-affinity self-reactive B cells aberrantly survive,^7, 8^ and glucocorticoid-induced leucine zipper (GILZ)-deficient mice^9^ with impaired regulation of activated B cells. These and various other mouse models of SLE replicate some aspects of disease relevant to some patients with SLE, but most likely do not individually account for all the disease symptoms and pathogenesis mechanisms in humans.

Numerous large-scale clinical trials for SLE treatments have been carried out, with an improvement over standard of care as the expected outcome of these studies. Disappointingly, the vast majority of tested therapies failed their primary endpoints,^10^ except belimumab, an inhibitor of the cytokine BAFF, showing modest efficacy in a subset of patients with SLE.^11^ Highly variable responses to treatments could be explained by the fact that recruitment of patients into clinical trials is based on a limited set of clinical manifestations and/or clinical scores, unlikely to fully capture the differences between patients. Therefore, there is an unmet need for more meaningful patient stratification and recruitment criteria, not just limited to clinical manifestations. Indeed, this can be better achieved using biomarkers reflecting the specific underlying mechanism of disease, allowing for a more mechanism-targeted and personalised approach to therapy.

Here, we have applied machine learning approaches to stratify patients with SLE based on gene expression patterns derived from whole-blood transcriptome data. We demonstrated that this approach can better harness disease heterogeneity than clinical observations alone and can identify patient clusters with different biological mechanisms underpinning disease.

## MATERIALS AND METHODS

### Human subjects

Human subjects in Datasets 1 and 3 are previously described (table 1).^12, 13^ Patients with SLE and healthy donors in Dataset 2 were recruited from the Monash Medical Centre.^14^ Patients with SLE fulfilled the American College of Rheumatology (ACR) classification criteria.^15^ The SLE disease activity index 2000 (SLEDAI-2k)^16^ and the Physician Global Assessment (PGA; range 0-3)^17^ scores were recorded. Blood was collected into PAXgene Blood RNA tubes (BD), which were frozen at −20 °C for later RNA extraction. Patients did not participate in the analysis.

**Table 1.** Cohorts of patients and healthy donors, for whole-blood RNA-seq data.

| Dataset & Reference | Subjects | Collection Site | Clinical Metadata | RNA Sequencing Method |
| --- | --- | --- | --- | --- |
| **Dataset 1** |  |  |  |  |
| **Hung *et al.* (2015)<sup>12</sup>** | 99 SLE (93 female, 6 male). | UCSF Medical Center, USA. | • Anti-Ro ('none', 'medium', 'high')<br>• ISM ('low', 'high') | • Whole-blood collected in PAXgene tubes, RNA extracted with TRIZOL (Ambion).<br>• RIN checked but not specified.<br>• TruSeq library preparation kit (Illumina).<br>• HiSeq2000 platform (Illumina).<br>• 50 bp SE reads. |
| Accession: PRJNA294187 | 18 healthy (female). |  |  |  |
| **Dataset 2** |  |  |  |  |
| **This study.** | 30 SLE (28 female, 2 male). | Monash Medical Centre, Melbourne, Australia. | • Age<br>• SLEDAI-2k, PGA<br>• Clinical manifestations<br>• Flow cytometry<br>• Medications | • Whole-blood collected in PAXgene tubes, RNA extracted with PAXgene kit.<br>• RIN > 7.<br>• TruSeq library preparation kit (Illumina).<br>• HiSeq 2500 platform (Illumina).<br>• 100 bp SE reads. |
| Accession: PRJNA439269 | 29 healthy (27 female, 2 male). |  |  |  |
| **Dataset 3** |  |  |  |  |
| **Rai *et al.* (2016)<sup>13</sup>** | 12 patients with SLE. | Sir Sunderlal Hospital, Banaras Hindu University, India. | • Age<br>• SLEDAI-2k<br>• Anti-DNA (±)<br>• Anti-ENA (±)<br>• Clinical manifestations<br>• Medications | • Whole-blood collected in heparin tubes, RBC lysis buffer, RNA extracted with TRI reagent (Sigma).<br>• RIN > 7.<br>• TruSeq library preparation kit (Illumina).<br>• HiSeq2000 platform (Illumina).<br>• 100 bp PE reads. |
| Accession: PRJNA318253 | 4 healthy donors. |  |  |  |
|  | All female. |  |  |  |
| **Meta-analysis** |  |  |  |  |
| **This study.** | 141 SLE | As above. | As above. | As above. |
| Datasets 1+2+3. | 51 healthy |  |  |  |

All RNA-seq data are publicly available from the Sequence Read Archive (SRA).<sup>59</sup>
Excluded sample in Dataset 2: "SLE\_21" (SRR6970317) was later found to not have SLE.
Abbreviations: ENA, extractable nuclear antigens; ISM, interferon signature metric; PE, paired-end; PGA, Physician Global Assessment; RIN, RNA integrity number; SE, single-end; SLE, systemic lupus erythematosus; SLEDAI-2k, SLE disease activity index 2000; UCSF, University of California San Francisco.

### RNA extraction and RNA-sequencing

RNA was extracted using PAXgene Blood RNA kits (Qiagen). RNA libraries were prepared for sequencing using standard Illumina protocols. RNA-sequencing (RNA-seq) was performed on an Illumina HiSeq 2500 platform; 100 bp single-end, stranded reads were analysed with the bcl2fastq 1.8.4 pipeline. Sequence read data is available on Gene Expression Omnibus (GSE112087).

### Bioinformatics analysis

#### Read quality, trimming, mapping, and summarisation

Publicly available datasets used in this study are listed in Table 1.^12, 13^ RNA-seq data was processed using a consistent workflow (supplementary figure S1). All software is listed in supplementary table S1. Read ends were trimmed with Trimmomatic (v0.38) using a sliding window quality filter.^18^ Datasets 2 and 3 were truncated to 50 bp single-end format to be consistent with Dataset 1, before read mapping. Reads were mapped using HISAT2^19^ (v2.1.0) to the human reference genome GRCh38/hg38 and the GENCODE Release V27 of the human genome GRCh38.p10 was used to annotate genes (35,398 genes included). Read counts were summarised using the *featureCounts* function of the Subread software package (v1.6.1);^20^ non-uniquely mapped reads (i.e. reads which map to more than one gene ambiguously) were excluded from analysis. Males (10% of patients) were included but Y-chromosome genes were excluded from the analyses. Lowly expressed genes were filtered out using a threshold requiring at least 0.5 counts per million (cpm) in healthy donor samples across all datasets. In total, 9,983 genes with unique Entrez accession numbers were retained.

#### Normalisation and standardisation

Read counts were normalised by the upper-quartile method, to correct for differences in sequencing depth between samples, using edgeR.^21, 22^ Counts were log2 transformed with an offset of 1, and samples in each dataset were computed as the log2 fold-change (log_2_FC) against the matching healthy control group mean. These processing steps were useful to reduce the distracting effects of extreme values and skewness typically found in RNA-seq data.^23^

#### Gene selection, clustering, and machine learning

Principal components analysis (PCA) and sparse partial least squares discriminant analysis (sPLS-DA) was performed using the mixOmics R package (using Lasso penalisation to rank predictive genes),^24^ and the MUVR R package (v.0.0.971).^25^ Cross-validation was used to protect against overfitting: in mixOmics, using M-fold cross-validation (10-folds averaged 50 times); in MUVR, using 15 repetitions of repeated double cross-validation (rdCV). A repeated measures design was used when combining datasets.^26^ Unsupervised clustering was performed with MATLAB (MathWorks), using the *k*-means function (using 100 repetitions to optimise initial centroid positions). Error-correcting output codes (ECOC) classifiers, which contain several support vector machines for multi-class identification, were generated using MATLAB. Random forest classifiers were generated using MUVR.^25^

### Differential gene expression and gene set enrichment analysis

#### Count-based expression analyses

The limma/edgeR workflow was used for differential expression analysis.^22^ The EGSEA (v1.10.1) R package was used to statistically test for enrichment of gene expression sets, using a consensus of several gene set enrichment analysis tools.^27^ EGSEA uses count data transformed with *voom* (a function of the limma package).^28^ Collections of pre-defined gene sets were from KEGG Pathways, and the Molecular Signatures Database (MSigDB: “H”, “c2”, and “c5” collections).^29^

### Circulating immune cell composition analysis

#### Flow cytometry

Whole blood samples collected into lithium heparin tubes (BD) were examined for frequency of circulating neutrophils (CD16^+^, CD49d^−^) by flow cytometry, using an LSR Fortessa instrument (BD Biosciences), and FlowJo software (Tree Star), as previously described.^30^

#### Transcript-length-adjusted expression and cell type enrichment analysis

Transcript-length-adjusted expression estimates (FPKM, or Fragments Per Kilobase of transcript per Million mapped reads) were obtained using StringTie (v1.3.4) and Ballgown (v2.12.0) R packages.^19^ Whole-blood RNA-seq results (FPKM format) were analysed for immune cell type signature enrichment using the xCell R package (v1.1.0).^31^

### Statistical Analysis

The mixOmics and MUVR R packages were used for multivariate analysis using count data.^32^ R version 3.5.2 was used. Kruskal-Wallis tests (with Dunn’s correction for multiple comparisons) and Mann-Whitney tests were performed using Prism software (v8.0.2, GraphPad). Statistically significant differences are shown for p < 0.05 (*), p < 0.01 (**), p < 0.001 (***); p < 0.0001 (****); or not significant (n.s.).

## RESULTS

We examined our cohort of 30 patients with SLE and 29 healthy donors for differentially expressed genes by RNA-seq, alongside two other publicly available datasets (141 SLE and 51 healthy donor whole-blood transcriptomes in total). Principal components analysis (PCA), which looks at all gene expression and visualises the overall variance between individuals, suggests a higher gene expression heterogeneity in SLE samples than healthy controls, which projected more closely together (figure 1A). Gene expression in some SLE samples was similar to that of healthy controls. Supervised clustering (to draw apart the groups) was performed using sparse partial least squares discriminant analysis (sPLS-DA). This method selected a subset of discriminating genes that are more useful for separating healthy and SLE patients (figure 1B). An expression heatmap using the top-ranking discriminating genes shows heterogeneity across patients with SLE (figure 1C), but visually demonstrates the possibility of organising SLE patients into several discrete clusters.

**Figure 1.**
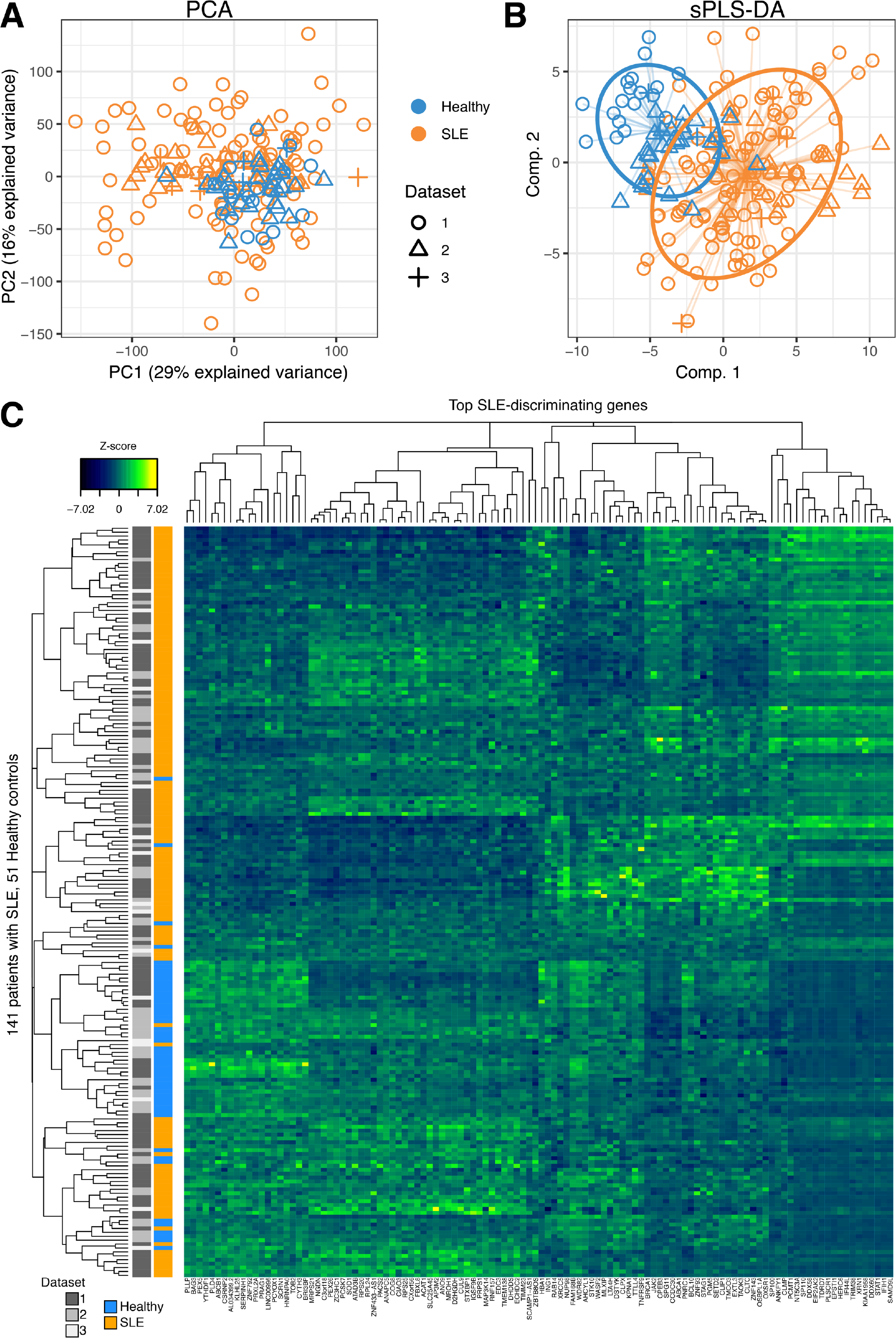
Differential gene expression in SLE. 141 SLE (orange symbols) and 51 healthy donor (blue symbols) transcriptomes from three datasets (see table 1, shown with different symbol shapes), were examined using multivariate statistics methods. (**A**) Principal coordinates analysis (PCA) was applied to visualise the overall variance between individuals. (**B**) Sparse partial least squares discriminant analysis (sPLS-DA), a supervised clustering method, applies weighting to genes which separate healthy donors and unstratified SLE patients. Ovals indicate the 80% prediction interval. (**C**) Standardised expression of top-weighted genes from the sPLS-DA model were plotted as a heatmap. Each row is an individual, each column is a gene.

We applied unsupervised *k*-means clustering to group patients into four clusters, C1-C4; Clusters were visualised with a PCA plot (figure 2A). The *k*-means clustering algorithm uses a chosen number of cluster centroids, which are repositioned among the samples until convergence.^33^ Supervised machine learning was applied, confirming that classification software can be trained to learn the transcriptomic signatures of each cluster and accurately classify new patients (88% accuracy, supplementary figures S2-3, using two different classifier algorithms).

**Figure 2.**
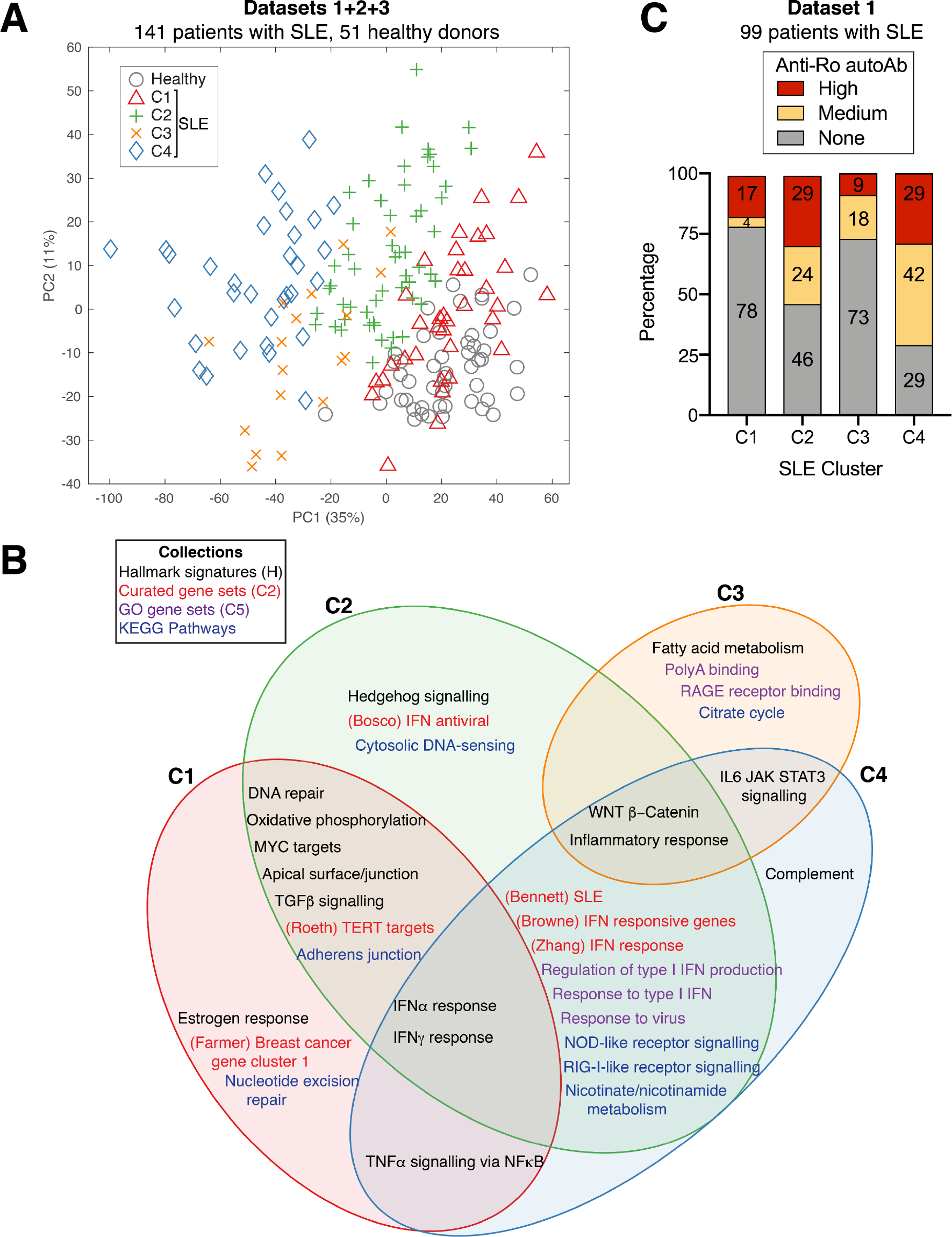
Patient clustering. (**A**) PCA visualisation of 141 SLE whole-blood transcriptomes after clustering using the *k*-means algorithm. Four clusters of patients were segregated and displayed with different symbols. Three datasets were combined (see Table 1). (**B**) Venn diagram displaying selected top-ranking disturbed gene sets in each SLE cluster C1-C4 compared to the healthy control group (derived from 99 patients with SLE and 18 healthy donors from Dataset 1). (**C**) Percentage of anti-Ro autoantibody levels in 99 patients from Dataset 1, rated as “none”, “medium” or “high”, derived from Dataset 1 metadata.^12^

Cluster 1 (C1) is transcriptionally the most similar to healthy donors, compared to C2-C4, which have incrementally more differentially expressed genes (supplementary figure S4). Gene set enrichment analysis was performed to summarise the predominant transcriptomic differences between the clusters (figure 2B). The top-ranking disturbed pathways, which differentiate the clusters, include immune activation pathways (e.g. anti-viral interferon response), metabolic pathways (e.g. citrate cycle), and DNA repair gene sets. Some of the pathways are likely attributable to particular medications, such as reactive oxygen species (ROS) generation gene sets, which are a known effect of hydroxychloroquine treatment.^34^

Interestingly, anti-Ro autoantibody positivity was substantially increased in C2 and C4 (figure 2C). Ascending levels of overall disease severity were observed from cluster 1 to 4, as suggested by the SLEDAI-2k (figure 3A), and PGA scores (figure 3B). Anti-dsDNA autoantibody ratio is significantly increased in C4 compared to the other clusters (figure 3C).

**Figure 3.**
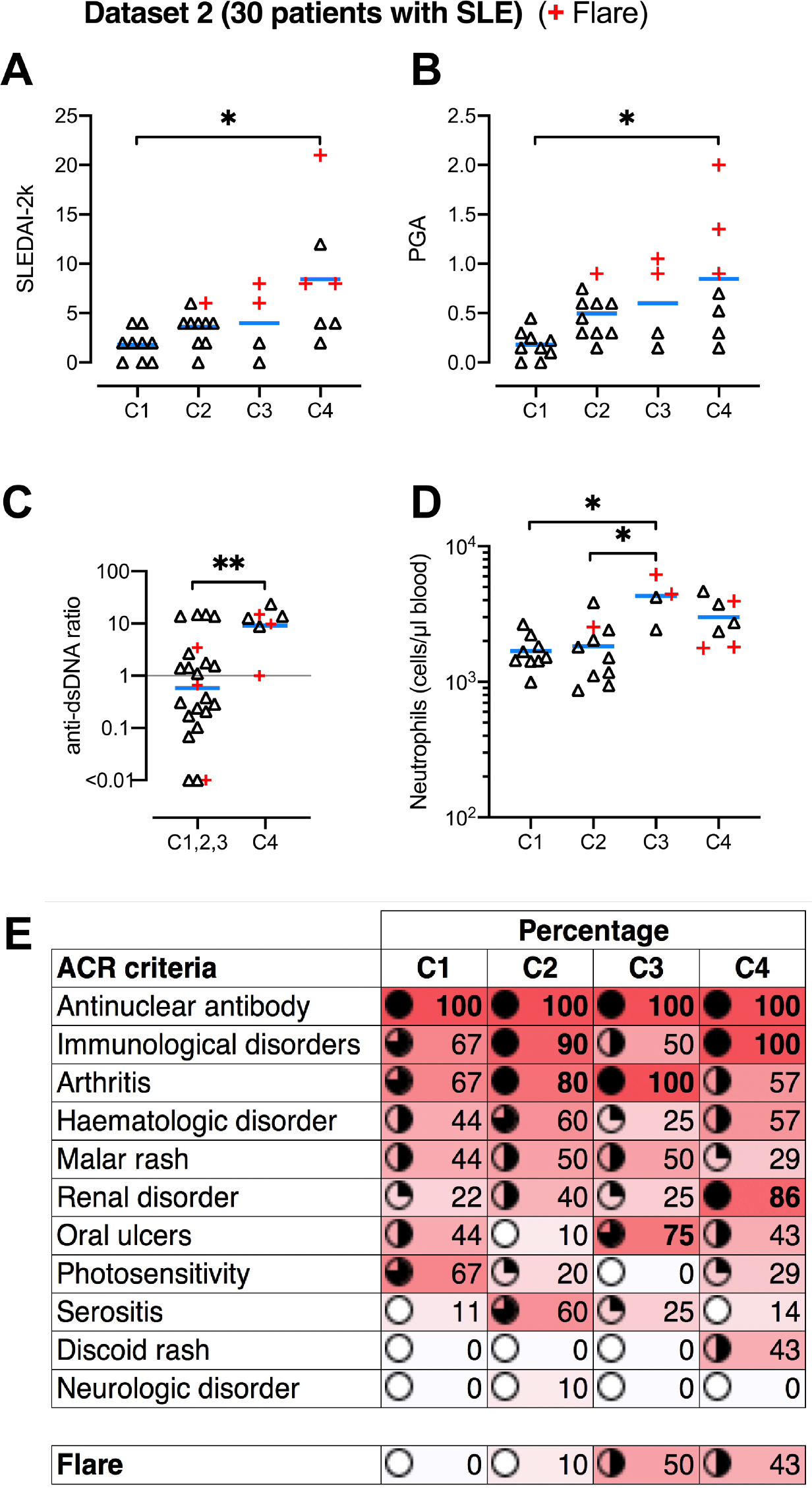
Disease severity and clinical features in SLE subtypes. SLE clusters C1-C4 in Dataset 2 were compared by clinical features. Blue bars represent the mean, symbols represent patients. Red (+) symbols represent patients experiencing flares (temporary period of worsened symptoms) at the time of sampling. (**A**) SLE disease activity index 2000 (SLEDAI-2k). (**B**) Physician Global Assessment (PGA). (**C**) Ratio of anti-dsDNA autoantibodies, in C4 vs the other clusters combined. (**D**) Circulating neutrophil numbers. (**E**) Percentage map of patients in each cluster, who are positive for particular disease features as detailed (ACR criteria) and flare activity.

Flow cytometry revealed that circulating neutrophil numbers were significantly increased in C3 (figure 3D). “xCell” (a software tool looking at cell-specific genes) ^31^ calculated enrichment scores, suggesting several cell-type compositions differences (supplementary figure S3). In particular, plasma cell gene signature is reduced in C3, B cell and CD8^+^ T cell gene signatures are reduced in C3 and C4; NKT cell gene signature is increased in C4, while that of conventional dendritic cells (cDC) is reduced in C4. M1 and M2 macrophage gene signatures are not significantly altered (supplementary figure S3).

The 30 patients in Dataset 2 all presented with a similar total number of American College of Rheumatology (ACR) criteria, although there are marked differences in the type of ACR criteria in each cluster. For instance, C4 has the highest positivity for immunologic/renal disorders and flare activity (suggesting more serious disease severity), whereas C2 and C3 have the highest positivity for arthritis (figure 3E).

In comparing the expression levels of several well-established SLE-associated genes in SLE clusters, we found evidence that different pathogenesis pathways were associated with each cluster of patients. BAFF (*TNFSF13B*) overexpression is well-established as a driver of autoimmunity,^7^ targeted by belimumab. Interestingly, high BAFF expression is a very significant feature of C4 and to a lesser magnitude C2, but not C1 and C3 (figure 4A). *TNFSF10* mRNA (encoding the apoptosis-inducing ligand TRAIL) expression is also upregulated in SLE,^35^ and this mirrors elevated BAFF expression in C4 and C2 (figure 4B). Defective apoptosis has been implicated in autoinflammatory settings, including SLE.^36^ Efficient apoptosis can be impaired by upregulation of anti-apoptotic factors such as cellular FLICE-inhibitory protein (encoded by *CFLAR*), previously reported to be upregulated in blood B cells of patients with SLE, and correlating with disease severity.^36^ This likely prevents apoptosis signalling in response to ligands such as TRAIL and FASL, to allow aberrant survival of autoreactive cells.^36^ Our stratification found substantial *CFLAR* overexpression in C3 and C4 (figure 4C).

**Figure 4.**
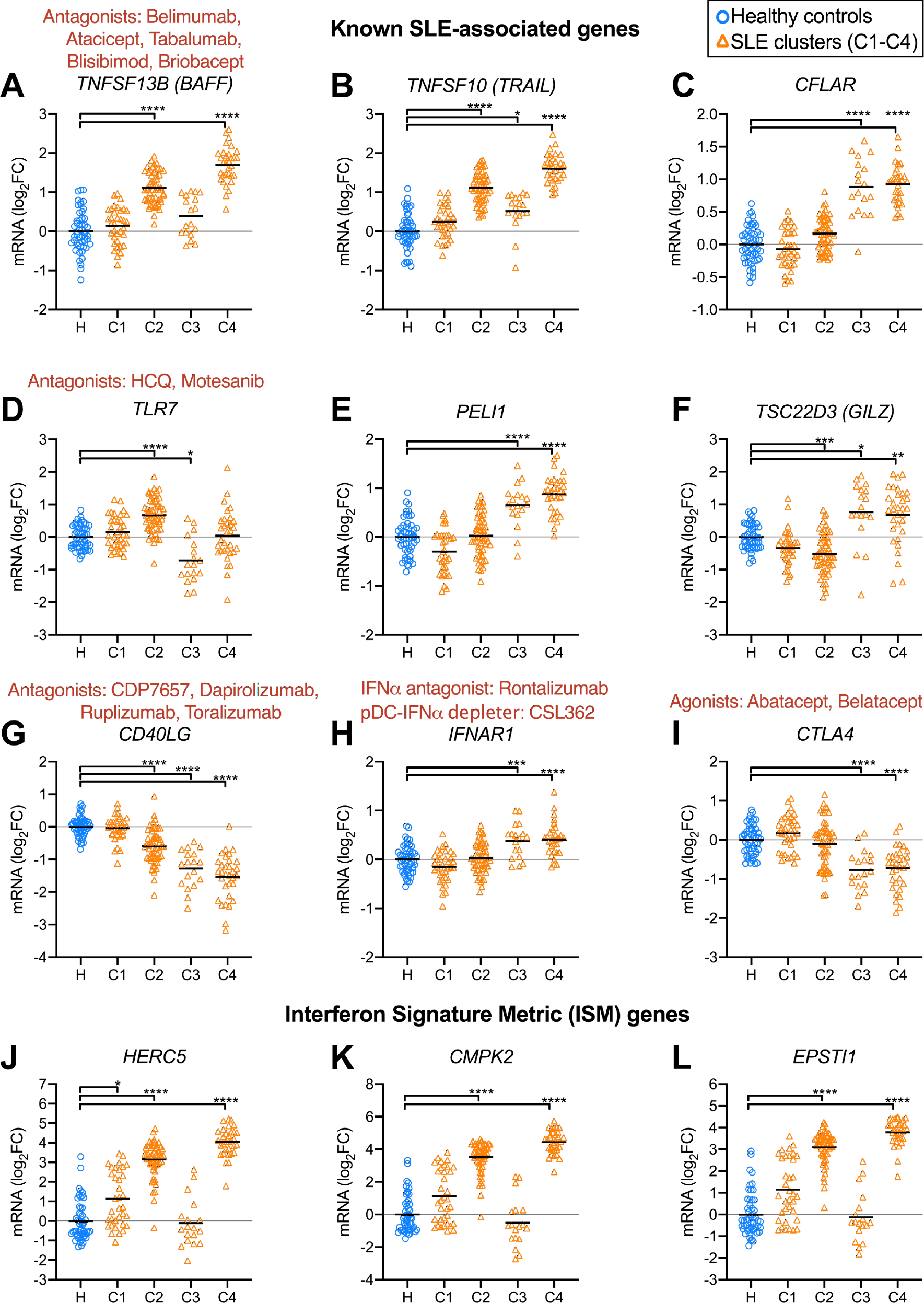
Relative expression levels of known SLE-associated genes. Expression levels (Log2 fold-change relative to the mean of the healthy controls) of (**A**) *TNFSF13B* (*BAFF*), (**B**)*TNFSF10* (*TRAIL*), (**C**) *CFLAR*, (**D**) *TLR7*, (**E**) *PELI1*, (**F**) *TSC22D3* (*GILZ*), (**G**) *CD40LG*, (**H**) *IFNAR1*, (**I**) *CTLA4*. Expression of interferon signature metric (ISM) genes: (**J**) *HERC5*, (**K**)*CMPK2*, and (**L**) *EPSTI1*. Therapeutics are indicated in red text above genes coding for the relevant target protein. Three datasets were combined (see Table 1). Significant differences (detailed in methods) between healthy and SLE samples are indicated.

Excessive TLR receptor signalling is implicated in autoimmunity, with TLR2, TLR7 and TLR9 pursued as potential therapeutic targets in SLE.^37^ Deregulated excessive TLR signalling is thought to exacerbate unspecific immune cell activation.^38^ Interestingly, TLR7 expression was significantly upregulated in C2 and downregulated in C3 (figure 4D). *PELI1* (encoding Pellino1) is a TLR3-inducible negative regulator of noncanonical NF-κB and the expression of *PELI1* was negatively correlated with disease severity.^39, 40^ In our stratification, *PELI1* is not significantly under-expressed in any SLE clusters, but is upregulated in C3 and C4, possibly induced for NF-κB regulation (figure 4E). *TSC22D3* (also known as *GILZ*) was identified as a negative regulator of B cells and lack of *GILZ* can drive autoimmune disease.^9^ *GILZ* expression was markedly diminished in C2, suggesting a loss of B cell regulation. *GILZ* was upregulated in C3 and C4, possibly as an effect of glucocorticoid induction (figure 4E).

CD40L, encoded by *CD40LG*, mediates T-cell help driving T-dependent B-cell activation, and has been targeted in unsuccessful clinical trials.^10^ *CD40LG* expression was significantly diminished in clusters C2, C3, and C4, possibly reducing the usefulness of CD40L blockade in those patients (figure 4F).

*IFNAR1* expression is significantly increased in clusters C3 and C4, suggesting increased interferon signalling sensitivity (figure 3H). *CTLA4* expression is significantly reduced in C3 and C4, suggesting impaired regulation of effector T cells (figure 3I). The Interferon Signature Metric (ISM) is a composite score of mRNA expression from three interferon-regulated genes (*HERC5*, *CMPK2*, and *EPSTI1*).^41^ Expression of these genes was consistently upregulated in C2 and C4, whereas C3 levels were comparable to that of healthy donors. Patients in C1 had variable levels with only the expression of *HERC5* but not the other ISM genes being significantly increased relative to healthy controls (figure 4G-I).

In Dataset 2, 6 of the 30 patients with SLE had flares, who diverged further from healthy donors when visualised by PCA (figure 5A). Using sPLS-DA to select flare-discriminating genes (figure 5B), we found differential gene expression during flares to be consistent with increased innate activation and altered immune cell regulation (figure 5C-F). Indeed, the *RETN* gene, encoding the proinflammatory adipokine Resistin, is upregulated in patients with active flares only (figure 5C). Resistin is linked to proinflammatory cytokine induction.^42^ Significant downregulation of *TCL1A* and *PAX5* (figure 5D-E) during flares suggests alterations in T- and B-cell homeostasis, respectively.^43, 44^ *LCN2* expression is increased in patients with flares (figure 5F). *LCN2* encodes neutrophil gelatinase-associated lipocalin (NGAL), which suggests increased neutrophil-mediated anti-bacterial activity; NGAL is also a biomarker of kidney injury.^45^ Gene set enrichment analysis revealed a number of pro-inflammatory gene sets and a neutrophil gene signature are predominant features of flare activity (figure 5G).

**Figure 5.**
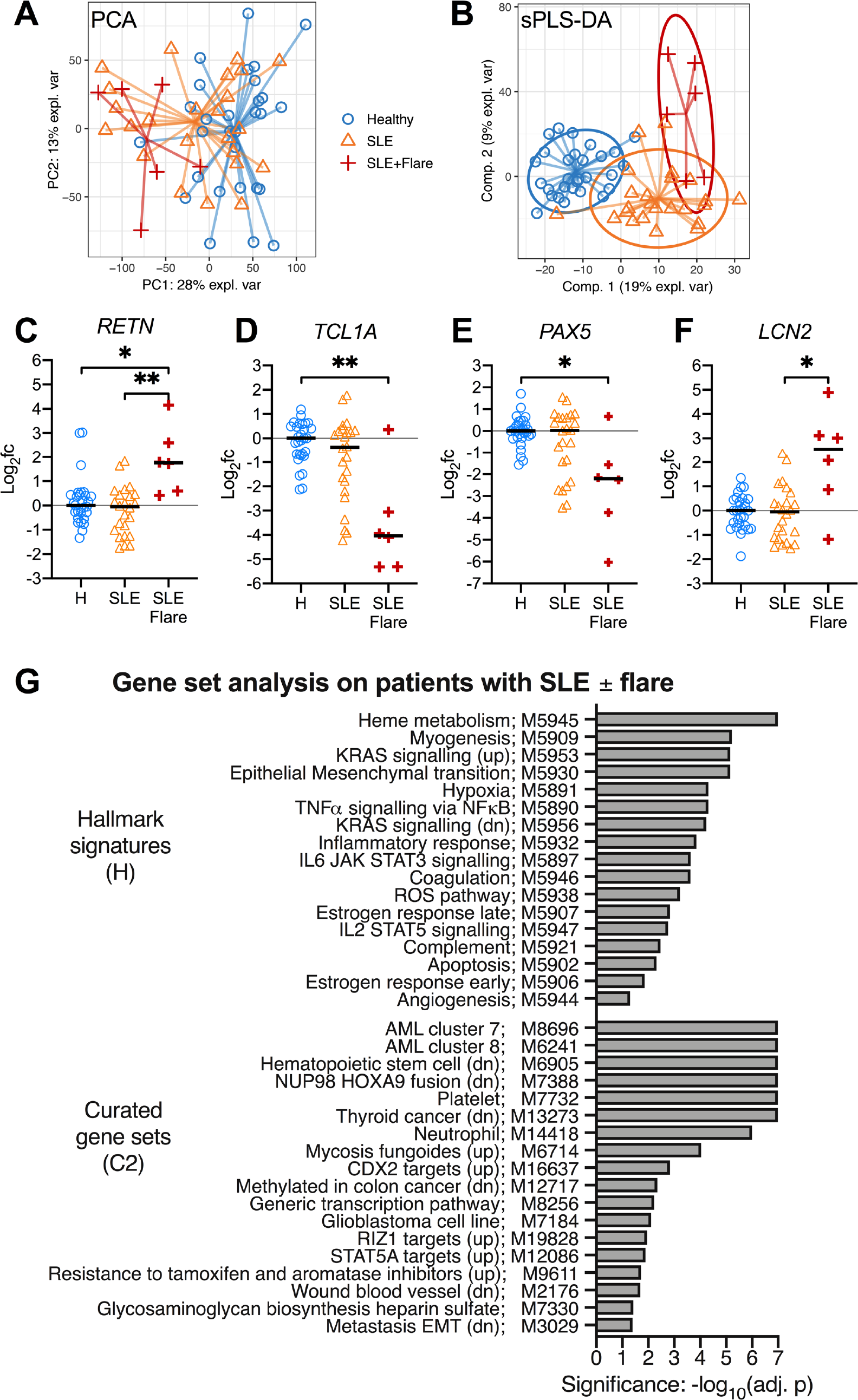
Gene signature for SLE flare activity. Whole-blood RNA-seq data from 30 SLE patients (24 without flares, and 6 with flares) and 29 healthy donors were compared (Dataset 2, see Table 1). (**A**) Principal components analysis (PCA) to visualise the variation between samples (in all genes); different symbols represent individuals in each group as shown. (**B**) Sparse partial least squares discriminant analysis (sPLS-DA) was used to select genes which distinguish the groups. (**C-F**) Relative expression of flare-associated genes, shown as the log2 fold-change relative to the mean of the healthy donor group (“H”) groups as shown. (**G**) Gene set enrichment analysis, showing the top-ranked gene sets which are differently expressed in patients with flares compared to patients without flares.

## DISCUSSION

A universally effective and safe treatment for SLE remains an unmet need due to the heterogeneity of clinical presentation, leading to an unpredictable response to treatment.^46^ SLE remains a condition with poor long-term outcome. Over six decades of clinical trials in SLE have only yielded one new therapy, belimumab, an inhibitor of the cytokine BAFF, with mixed efficacy in patients.^10^ Major failures of targeted therapy in the clinic for SLE^10, 47, 48^ mean that breakthrough treatments remain years away. This situation has obligated clinical experts and the pharmaceutical sector to more rigorously assess the reasons for this high failure rate. Suggested factors include issues with the design of clinical trials, difficulty in defining robust endpoints, non-ideal drug targets and biomarkers, and, high heterogeneity of study populations.^10^ Large-scale clinical trials invariably fail to demonstrate efficacy when enrolling patients selected on a limited number of clinical criteria, which do not capture the underlying molecular mechanism likely underpinning disease, which varies greatly in patients (figures 2-3). Inclusion of some patients with low disease propensity (C1) further weakens comparisons between placebo and experimental treatment groups.

Our stratification method captures the likely underlying disease mechanism, using whole-blood transcriptomics to obtain a snapshot of the immune system. This stratification could be very useful for the improved design of clinical trials, by more appropriately targeting specific clusters of patients with SLE who are much more likely to have a homogenous mechanism of action underpinning pathology (figures 2B,4). Retrospective analysis of previous failed trials could reveal high efficacy in specific clusters of patients, which was not possible to see in an unstratified analysis. Successful off-label usage of rituximab in some patients with SLE further suggests therapies that have failed in clinical trials with SLE may yet have efficacy in selected patients.^49, 50^ Indeed, looking at the expression levels of key drug-targeted molecules such as BAFF and CD40L suggests that certain clusters of patients might be much better fit for the rationale of targeted biologics than other clusters (figure 4).

Similar to us, previous studies using microarrays have encountered distinct clusters of SLE patients in whole-blood transcriptome data.^51, 52^ In this study, we used RNA-seq data, which has the advantages of capturing additional genes (not restricted to probe sets) and improved dynamic range. Additional systems biology approaches (such as microbial metagenomics, and metabolomics) are becoming available in SLE, and combining matching data from additional profiling methods may allow for improved sets of clinically useful biomarkers.^53–56^

Transient flare activity in SLE patients causes a significant surge in inflammation requiring medical attention, but much remains to be understood about the underlying molecular basis and transcriptomic features of flare activity. We identified several flare-associated genes including the *RETN* gene, encoding the proinflammatory adipokine resistin (figure 5C). Interestingly, serum resistin levels are elevated in patients with rheumatoid arthritis and/or SLE patients, although the differences were reported not significant in unstratified patients with SLE, where high heterogeneity was noted.^57^ The specificity of elevated resistin levels to flare-active patients may explain these results. Taking this further, longitudinal studies would be useful for discovering flare-predicting transcriptional signatures, which may be used as prognostic markers alerting patients and physicians of an increased risk of flares under their current treatment plan.

The IFN gene signature was associated with patients with SLE, although this feature does not correlate well with overall disease severity.^41^ Stratification of ISM-high patients is possible using qPCR assays for the gene expression of three genes in peripheral blood,^41^ which in our stratification corresponds to C2 and C4 (figure 2B, 4H-L). Several new treatments related to type I interferon are under investigation, for example, anti-IL3Ra (i.e. anti-CD123, CSL362 mAb), which depletes basophils and plasmacytoid dendritic cells, a cell type which produces type-I IFN.^30^

In conclusion, our study provides new insights into the heterogeneity of patients with SLE with respect to gene expression in circulating immune cells, as the messengers of overall immune activity in individual patients. Our approach using whole-blood transcriptomics data combined with machine learning approaches is powerful at segregating and recognising patient clusters, uncovering cluster-specific gene expression patterns linked to known pathogenesis features. Optimal patient stratification is critically lacking in clinical trials for SLE, for which success rates and cost-effectiveness can be greatly improved by more robustly targeting the most relevant clusters of patients. Further development of machine-learned classifiers and validation of their utility using matching data on patients’ response to specific therapies, may deliver new clinical tools assisting with better treatment decisions for individual patients.

## Acknowledgements

Computational work was performed using the high-performance computing (HPC) resources of the University of Melbourne^58^ (Project# punim0259), and Melbourne Bioinformatics (Project# UOM0044). We acknowledge the HPC training and technical assistance provided by the University of Melbourne, Melbourne Bioinformatics, and the Australian National Computational Infrastructure. This research was supported by use of the NeCTAR Research Cloud, a collaborative Australian research platform supported by the National Collaborative Research Infrastructure Strategy. We acknowledge Dr Kim-Anh Lê Cao for helpful discussions about multivariate statistics methods in the mixOmics R package.

## Authorship Contributions

WF conducted the analysis, wrote source code, produced the figures, and wrote the manuscript. FM, EFM, KM, MN, MA, and NJW reviewed the manuscript. KM, MN, MA, EFM, NJW, AH, and EFM generated Dataset 2.

## Funding

WF was supported by funding from the Victorian Cancer Agency (grant# ECSG15029).

## Competing interests

KM, MN, MA, EM, and NJW are employees of CSL Ltd.

## Patient consent

Obtained.

## Ethics approval

Blood.

## Supplementary Figures

**Figure S1.**
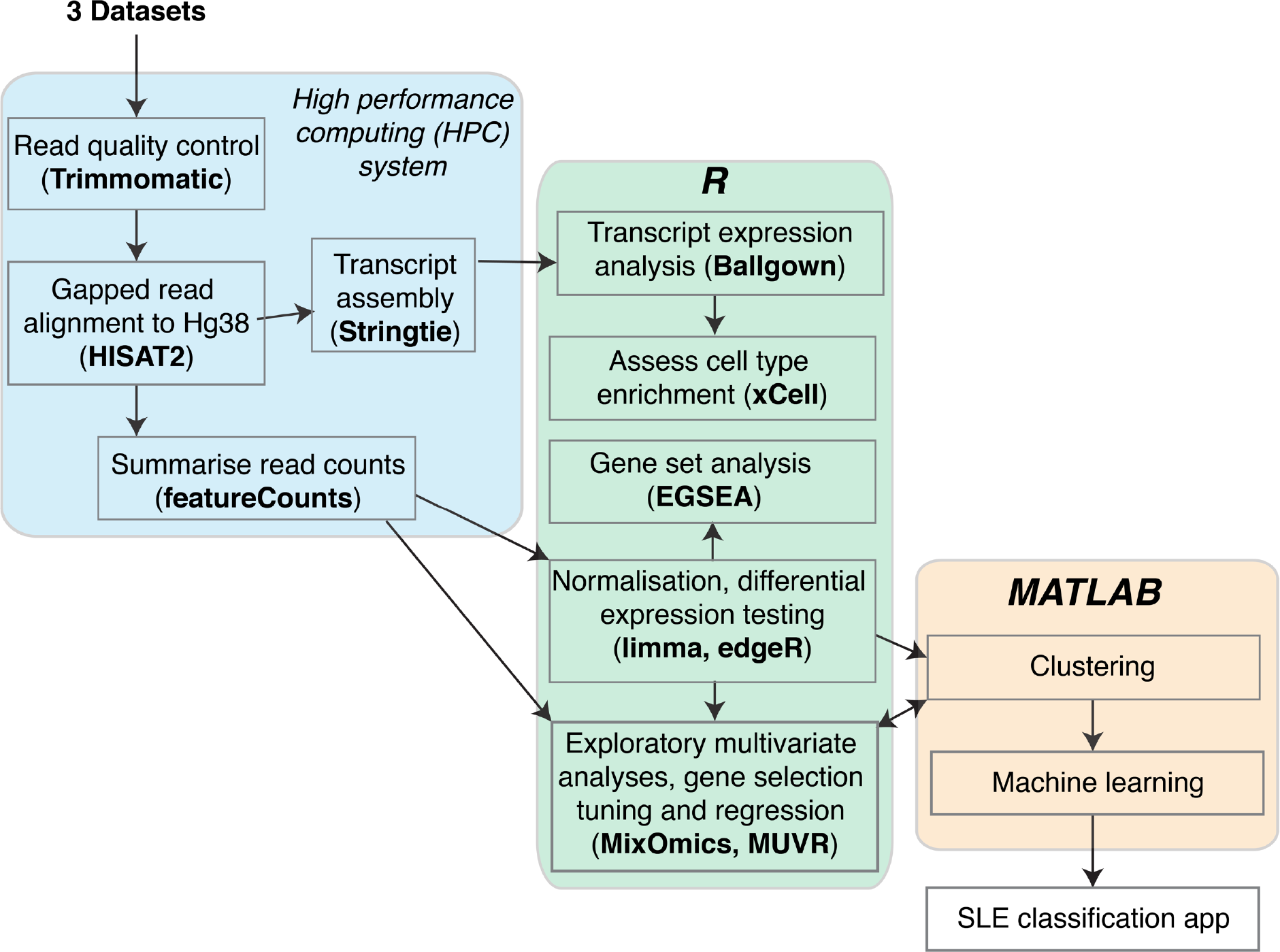
Bioinformatics workflow. Three RNA-seq datasets (supplementary table S1) were processed consistently using the depicted workflow.

**Figure S2.**
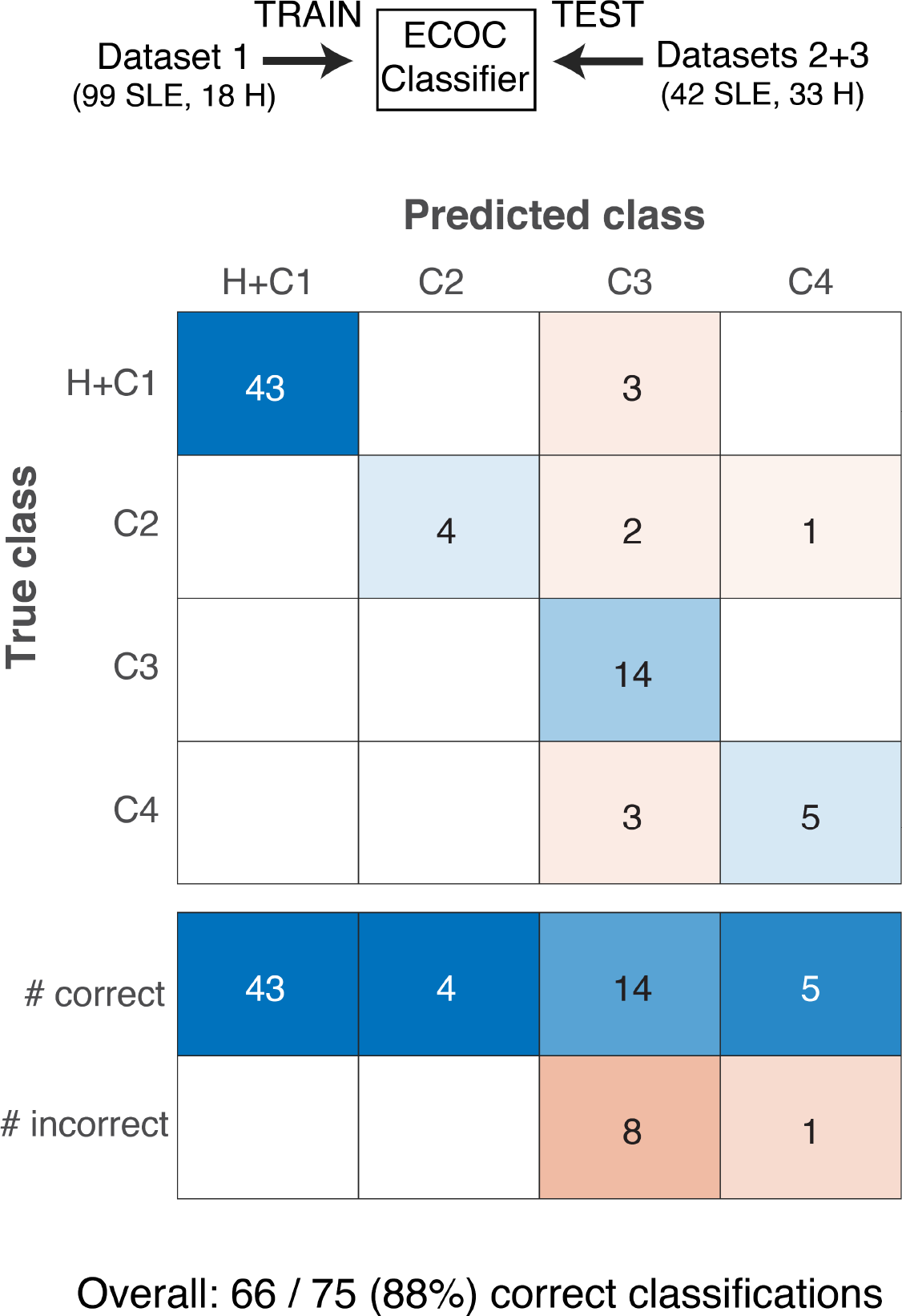
SLE subset discrimination using support vector machine classifiers. An error-correcting output codes (ECOC) classifier was trained using Dataset 1, to learn how to distinguish SLE clusters (C1-C4); in this case healthy donors were grouped with C1. The accuracy of the classifier was tested using independent cases from Datasets 2+3, checking whether the cluster identification matches the original clustering by *k*-means in figure 2.

**Figure S3.**
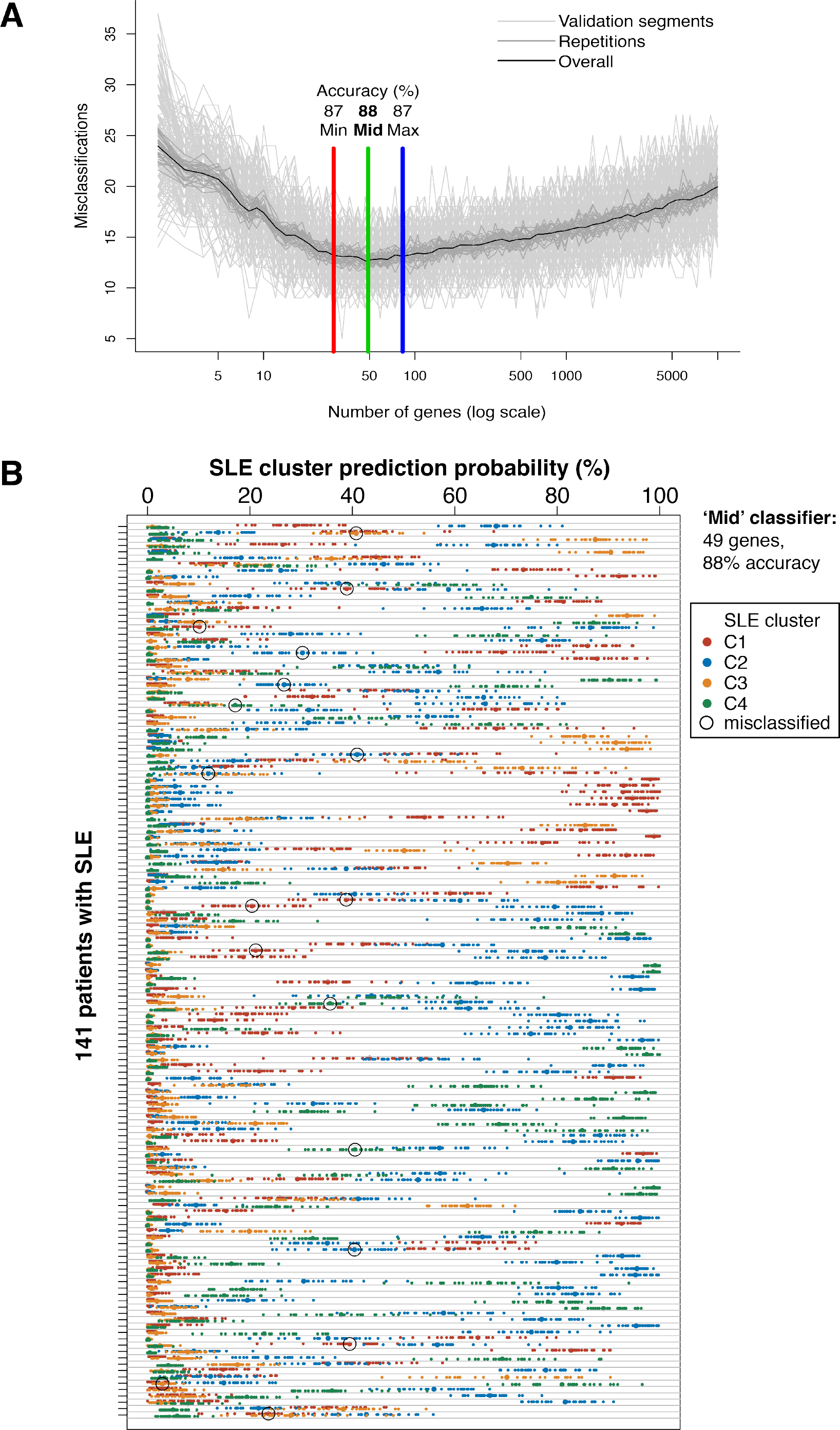
SLE subset discrimination using random forest classifiers. Three whole-blood RNA-seq datasets encompassing 141 patients with SLE were clustered (as in figure 1). (**A**) Random forest classifiers were trained and tested using repeated double cross validation to protect from overfitting, while selecting optimal gene sets with predictive value, using the minimum number of genes (’min’), maximum number of genes with predictive value (’max’), or the geometric mean from those models (’mid’). Using very few genes (on the left of this plot) results in a higher error rate; using too many genes with no added predictive value (on the right side of this plot) also results in a higher error rate due to the accumulation of noise. (**B**) Performance testing of the ‘mid’ classification model, which had 88% overall accuracy to predict the original cluster type, using 49 genes. For each sample (each horizontal lane), the predicted probability of each cluster designation (coloured symbols) is plotted. Incorrect classifications are circled. Smaller symbols show the result of test repetitions, larger symbols show the average result from the repetitions.

**Figure S4.**
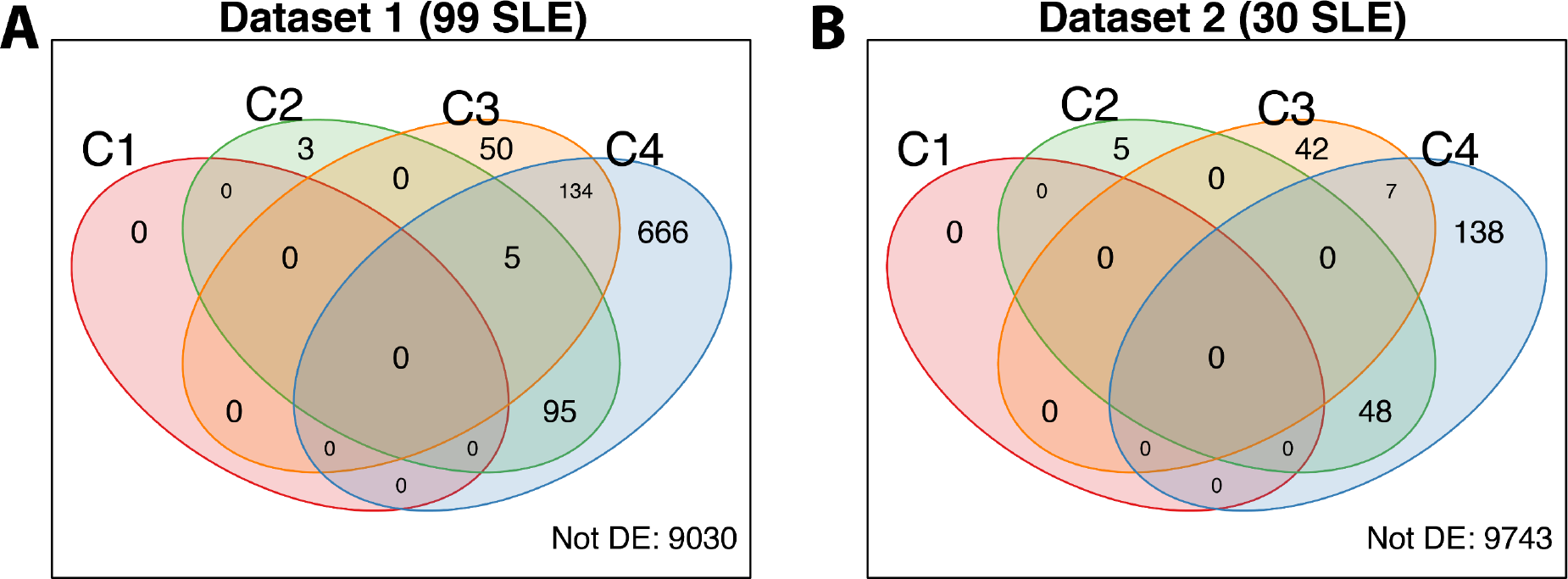
Differential Gene Expression in SLE clusters. Four SLE clusters (C1-C4) in Dataset 1 (**A**) and Dataset 2 (**B**) were analysed for differentially expressed (DE) genes compared to healthy donors, using the limma/edgeR workflow.^22^ The number of genes passing an arbitrary cut-off for differential expression (fold-change 1.5, BH-adjusted p-value 0.05) relative to Healthy donors are plotted as Venn diagrams. Using the same cut-off, fewer DE genes were found in Dataset 2 than Dataset 1 due to reduced cohort size, although the most DE genes were consistently found in C4 compared to other clusters regardless of dataset source.

**Figure S5.**
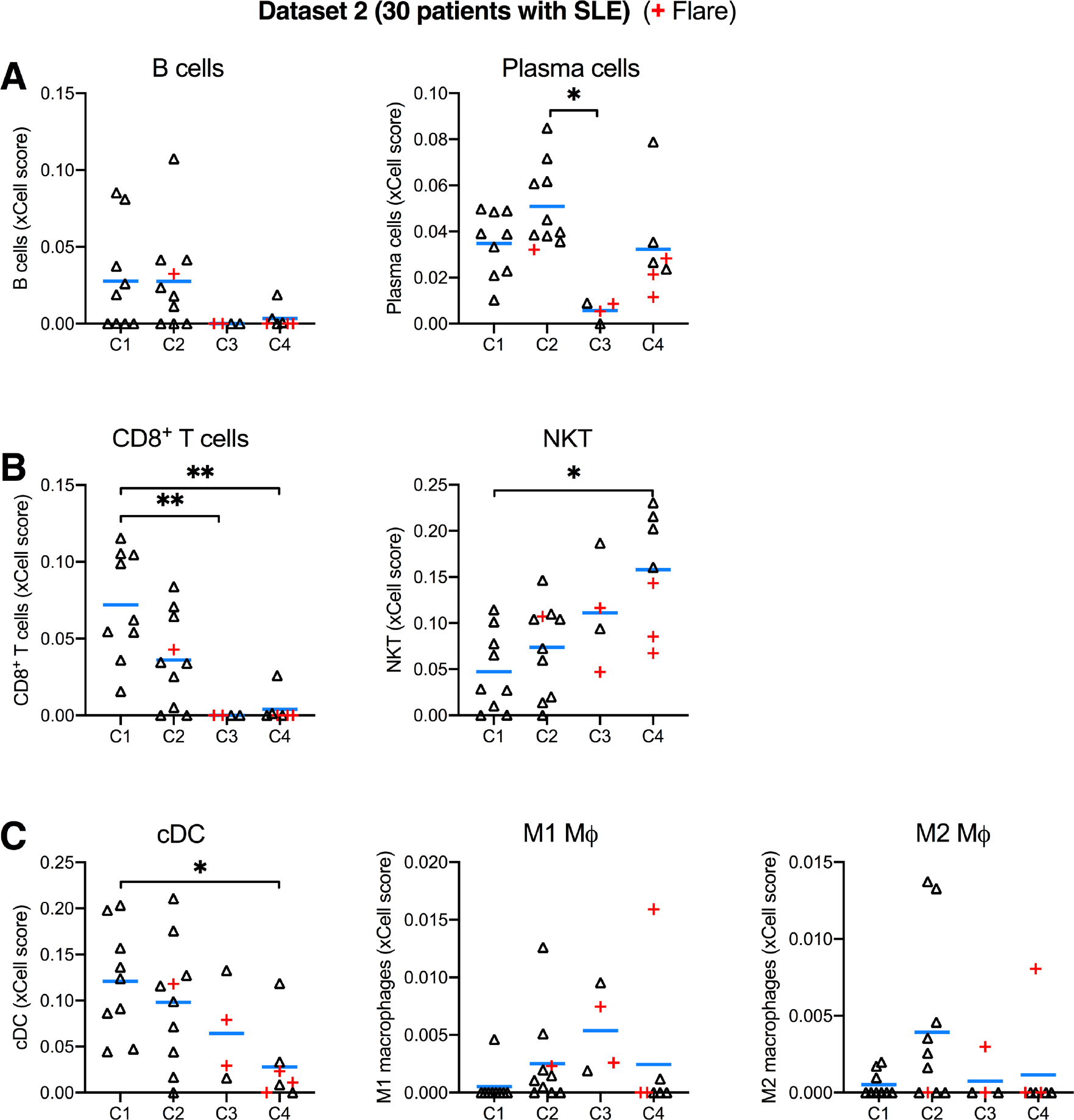
Cell subset deconvolution. 30 patients with SLE patients from Dataset 2 were stratified into four clusters (C1-C4); patients with flares are indicated with red (+) symbols. Blue bars show the mean. Immune cell type enrichment in whole-blood RNA-seq data was estimated from FPKM values using xCell.^31^ Signature enrichment scores for: (**A**) B cells and plasma cells; (**B**) CD8^+^ T cells, natural killer T cells (NKT); (**C**) conventional dendritic cells (cDC), M1 macrophages, M2 macrophages.

**Supplementary Table 1.** Software used for processing RNA-seq data.

| Software | Version | Purpose in this project | Ref. / Company |
| --- | --- | --- | --- |
| Ballgown | 2.12.0 | Calculate transcript abundance (FPKM). | 19 |
| edgeR | 3.24.3 | Count-based differential expression analysis. | 22 |
| EGSEA | 1.10.1 | Gene set enrichment analysis. | 27 |
| HISAT2 | 2.1.0 | Gapped read alignment. | 19 |
| limma | 3.38.3 | Count-based differential expression analysis. | 22 |
| MATLAB | 2018b | Clustering, machine learning. | MathWorks |
| mixOmics | 6.6.1 | Multivariate methods, variable selection. | 32 |
| MUVR | 0.0.971 | Variable selection, machine learning. | 25 |
| PRISM 8 | 8.0.2 | Graphing and statistical tests. | GraphPad Software |
| R | 3.5.2 | Statistical programming. | 60 |
| R Studio | 1.1.463 | Integrated development environment for R. | 61 |
| SAMtools | 1.8 | Sorting read alignments. | 62 |
| SRA-toolkit | 2.9.2 | *fastq-dump*: Obtain archived fastq data. | 59 |
| Stringtie | 1.3.5 | Transcript/splice model assembly. | 19 |
| Subread | 1.6.3 | *featureCounts*: summarise read counts at the gene level. | 20 |
| Trimmomatic | 0.38 | mRNA read trimming. | 18 |
| xCell | 1.1.0 | Cell-type enrichment analysis. | 31 |

